# Development and Implementation of a Minority Health International Infectious Diseases Research Training Program

**DOI:** 10.1101/2025.01.05.631114

**Authors:** Angela Sy, Diane Taylor, Pornsawan Leaungwutiwong, Nittaya Phanupak, Youngchim Sirida, Rose Leke, Peter S. Humphrey, Ravi Tandon, Madhur Kulkarni, Purnima Madhivanan, Joseph Keawe‵aimoku Kaholokula, Vivek R. Nerurkar

## Abstract

From 2014-2019, the University of Hawai‵i (UH) at Manoa offered a National Institutes of Health funded Minority Health International Research Training (MHIRT) Program for undergraduate research experiences in infectious diseases. The goal of the program was to immerse undergraduate students in conducting global infectious diseases research to train a new generation of scientists to combat future global pandemics. The MHIRT program trained educationally underrepresented groups unique in Hawai‵i: Native Hawai‵ians and other Pacific Islanders, and underrepresented Asian Americans, e.g., Filipinos and Vietnamese. A 12-member Advisory Committee guided the program development, implementation, recruitment strategies, recruitment, retention, and program evaluation. The program provided an eight-month classroom and field based international research training. Students learned about “8 steps of research” and cultural competency, experienced a hands-on research project abroad, and presented their results to the research community in the host country and at UH. Students’ research topics focused on ongoing projects of UH faculty including those with international collaborations in infectious disease and tropical medicine. Students spent two months on an international research mentor’s project at an international site under the mentorship of a Hawai‵i based research faculty. After eight months, students graduated from the program by presenting their research to family, friends, mentors, and UH faculty and administrators. Trainees in this program pursued academic graduate or professional health degrees. The majority of students pursued further studies related to infectious diseases contributing to workforce development in infectious disease health research. The UH-MHIRT program provides an example of student immersion in international infectious disease research to foster future interest in global health research providing potential benefits for the student and faculty and their research efforts.

---

Increases in the speed of global travel of humans and their goods have escalated the risk of infectious disease outbreaks, as evidenced by resurgence of the global coronavirus pandemic after a century since the previous global influenza virus pandemic. Newly emerging infectious diseases remain a priority in global health research perhaps now more than ever. The prevention and control of infectious diseases in international settings requires a complete understanding of the pathogens, disease manifestations and transmission mechanisms, ability to implement community preventive measures and interventions, and the extent to which health services and disease control are feasible (Baker et al., 2022).

Global health education in the United States has grown in demand because it provides experiential and engaged learning. The goals of the global health programs are to ultimately create a transformative and mutually beneficial learning for students, faculty, and affiliated partners (Mendes et al., 2020; Mosely et al., 2022). Study abroad programs provide students with the opportunity to integrate their classroom knowledge with applicable skills in an international setting (Kalbarczyk et al., 2015). Programs offering undergraduate research experiences often extend beyond traditional classroom and laboratory to challenge students to develop or identify appropriate researchable questions, experimental protocols, and analytical approaches to make sense of collected data (Olimpo et al, 2016). However, these programs are often offered locally within an academic institution. In these programs, institutions based in high-income countries offer short-term experiences in global health in low and middle-income settings which vary in length and purpose, including research training (Loh et al., 2015).

As academic global health training programs continue to grow, learning objectives for global health curricula and programs have been recommended by the Consortium of Universities for Global Health (CUGH 2024). Among global health learning objectives is the ability to specify “the burden of disease, disability, and death from infectious diseases” and ability to think “creatively about what policies and practices might prevent another emerging infectious disease pandemic” (Jacobsen, et al., 2020). Furthermore, World Health Organization Sustainable Development Goals 2030 indicates that the goals that are especially “stalled” in Africa are those that address, i) ending infectious diseases, ii) research and development of vaccines, and iii) medicine for diseases (Ending disease in Africa WHO, 2023).

National entities representing science and research have agendas to diversify the scientific workforce to include talented researchers from underrepresented groups to improve the nation’s capacity to address and eliminate health disparities. To address the compelling need to diversify the research workforce, the U.S. National Institutes of Health has developed grant programs to promote and support the training of racial and ethnic minorities and other populations underrepresented in biomedical, behavioral, clinical and social sciences research. The NIH-funded Minority Health International Research Training (MHIRT) T37 program provides both training experiences and the opportunity for innovative research collaborations and partnerships (National Institute on Minority Health and Health Disparities 2024).

The University of Hawai‵i at Manoa (UH Manoa), John A Burns School of Medicine (JABSOM) offers a MHIRT program drawing on formats and activities from study abroad programs, undergraduate research experiences, and global health education. Accordingly, program activities comprehensively train MHIRT students through coursework, laboratory, fieldwork, and community research experience abroad and domestically, and individual mentoring. Students are trained in scientific methods, laboratory techniques, analyzing and interpreting data, scientific literature reviews, scientific writing, responsible conduct of research, cultural competency, community engaged research and emerging international health issues. Students are also trained to address cultural, linguistic and ethical appropriateness and related issues while engaging in their biomedical research at the international site. Individual research mentoring as a training approach promotes students’ retention in their current academic degree programs and pursuit of advanced degrees in biomedical or behavioral health fields.

## Purpose

There is a need to diversify the research workforce by training underrepresented students in research especially in infectious diseases. MHIRT makes accessible for underrepresented students specific training in global infectious disease research.

### Objective

University of Hawai‵i-MHIRT (UH-MHIRT) goal is to increase the number of Native Hawai‵ian and other Pacific Islanders, Filipinos and other underrepresented students (rural, socio-economic) in biomedical research in Hawai‵i by providing a mentored international research experience in tropical medicine and infectious diseases in Asia and Africa. MHIRT fills important gaps in training that are not currently available in Hawai’i. The Department of Tropical Medicine, Medical Microbiology and Pharmacology (T3MP) programs at JABSOM aim to create *a pipeline of research trainees from high school through junior faculty*. However, as shown in Figure 1, the conspicuous gap in the pipeline at UH Manoa is undergraduate student training in infectious diseases. MHIRT was developed to fill this gap and help increase the number of underrepresented students in Hawai’i in biomedical research, especially those interested in tropical medicine and global health disparities (Figure 1). UH-MHIRT program is important because it lays out the pathway for educators in conducting global infectious diseases research training of significance for underrepresented students, to diversify the biomedical research workforce.

**FIG 1.**
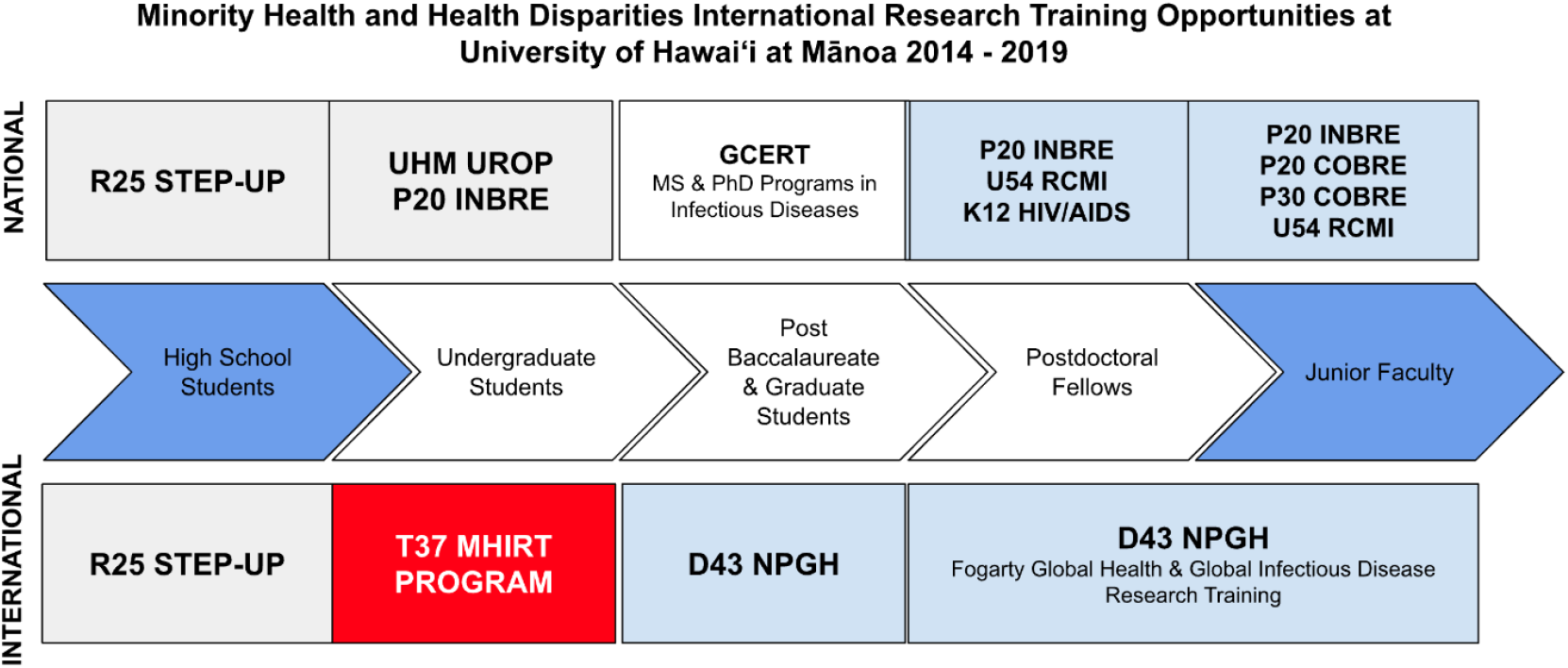
Minority Health and Health Disparities International Research Training Opportunities at the University of Hawai‵i at Manoa 2014-2019.

COBRE, Centers of Biomedical Research Excellence; GCERT, Graduate Certificate in Tropical Medicine; INBRE, IDeA Networks of Biomedical Research Excellence; NPGH, Northern Pacific Global Health; RCMI, Research Centers in Minority Institutions; STEP-UP, Short Term Research Experience Program to Unlock Potential; UROP, Undergraduate Research Opportunity

Hawai’i is geographically isolated, located ∼2,500 miles from the nearest USA landmass, and many underrepresented students have never left the state. Students from Hawai‵i are hesitant for economic, geographic, and cultural reasons to participate in MHIRT programs elsewhere in the continental US. Other MHIRT programs provide training for minority students, and only one program specifically has indigenous populations (Native Hawai‵ian, Alaska Natives, and American Indians (IWRI, 2024). The UH-MHIRT program is unique because it provides training for populations that are not served by other MHIRT programs in the U.S., i.e. Pacific Islanders and underrepresented Asians in biomedical research.

## Rationale

Conducting research in low- and middle-income countries (LMIC) is an eye-opening experience especially for students who have yet to leave the State of Hawai‵i, i.e., the Islands. The research collaborations at T3MP have seen first-hand health disparities in the diagnosis and treatment of tropical infectious diseases among people living in city and rural villages in Africa and Southeast Asia. MHIRT engages research trainees in infectious disease research to provide exposure to students to health disparities topics and to stimulate their interest in solving important (global) health problems. The Hawai‵i MHIRT program provides international research training in low-income (i.e., Liberia, Laos) and mid-income (i.e., Thailand, India, Cameroon, Palau) countries, creating a unique opportunity to compare problems related to health disparities in infectious diseases, tropical medicine, and other public health priorities (Figure 2). These countries were selected because the T3MP Department has long-standing international research project collaborations in these countries.

**FIG 2.**
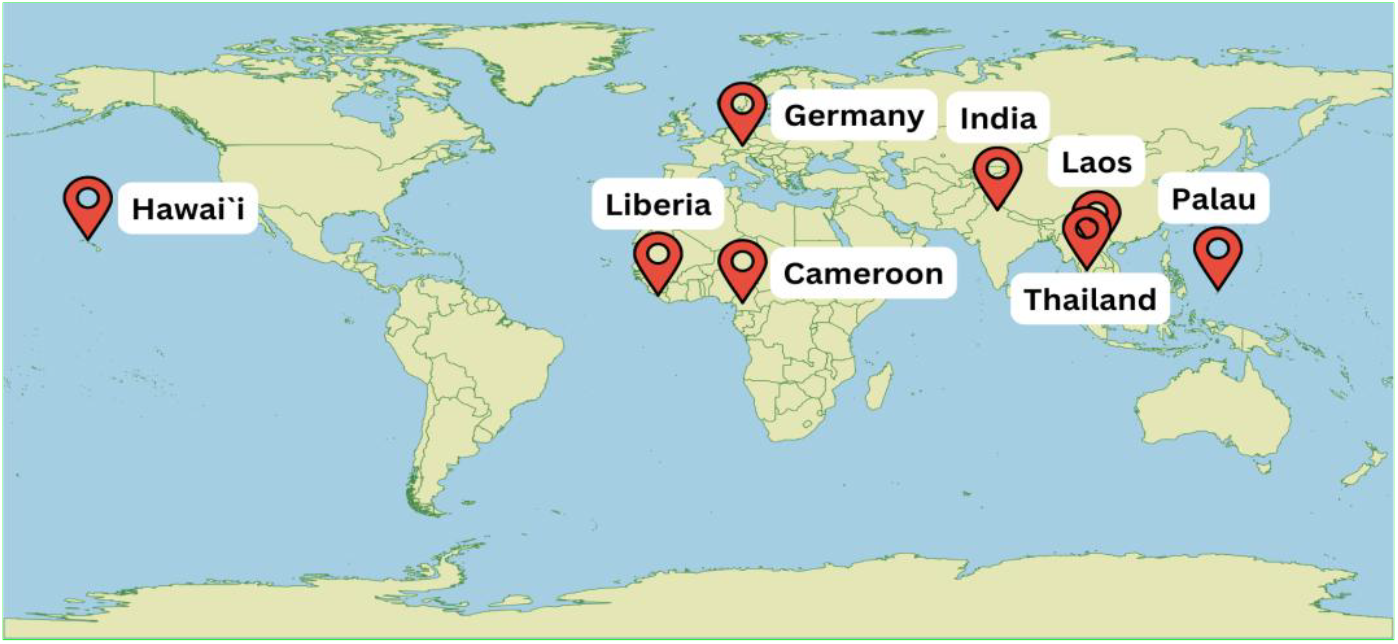
University of Hawai‵i international MHIRT sites, 2014-2019.

### Goals and Competencies

The MHIRT’s long term goal is to increase the future pool of global research scientists who are underrepresented including Native Hawai‵ians, other Pacific Islanders, Filipinos, and other categories of underrepresented students in Hawai‵i (e.g., rural, low-income). The goal is to annually engage and train nine underrepresented undergraduate and one graduate students in international tropical medicine research to instill an understanding of the role of biomedical research in helping overcome health disparities. Consistent with other programs to encourage and train underrepresented students in research, our program competencies focus on building student gains in research methods, responsible conduct in research, health disparities, and cultural humility (Supplement 1) (Ton, et al. 2015).

### Program Description

#### Overview of the Administrative Structure

The program has Co-Directors (PD) who administer the program with the support of a nine-member Advisory Committee. Seven core faculty at UH serve as academic advisors and research mentors for the students. Program site coordinators organize and oversee training in research laboratories at their foreign sites: Cameroon, Liberia, Thailand, India, Laos, Palau, and Germany (Figure 3). As needed, new training sites were added in the later years of the program. Trainees also have additional opportunities to conduct research throughout their academic careers at the UH.

**FIG 3.**
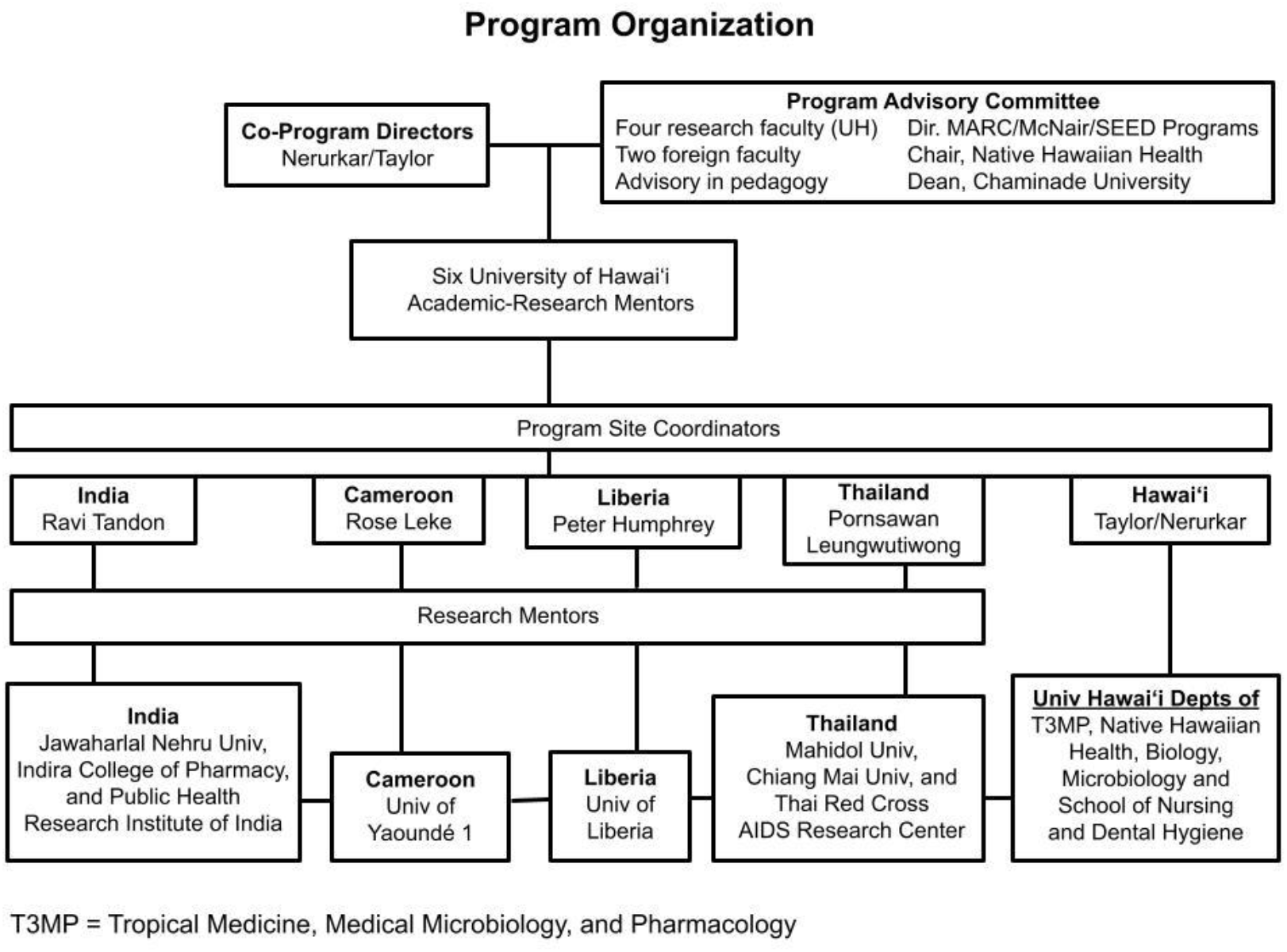
University of Hawai‵i MHIRT Program Organization Chart.

#### Advisory Committee

The voluntary Advisory Committee members’ duties and responsibilities were to assist in program development, implementation, recruitment strategies, recruitment, retention, and program evaluation. The Advisory Committee met once a year. The PDs, advisory committee members, and/or academic-research mentors from international sites met several times a year and in the interim if necessary to discuss problems as they arose, solutions and fine-tuning of the Program to optimize training of underrepresented students.

#### Research Mentors

For a successful international research training program, one needs mentors with diverse expertise in training undergraduate students. For UH-MHIRT, we had: i) Core academic-research mentors, ii) international mentors, and iii) on-site research mentors. Mentors are selected from infectious diseases and global health disciplines with teaching and research experience, including clinicians, with broad based knowledge. The role of these core mentors is to support students’ academic and professional interests and MHIRT program requirements. International mentors are accomplished scientists and strong student mentors. International mentors are from Thailand, India, Cameroon, Liberia, Laos, and Palau. They work locally with their institutions, and interactions and collaborations with Hawai‵i mentors are extensive including joint publications and R-series grant applications with U.S. researchers. UH research mentors are faculty already engaged in training minority students in research. Over 20 research faculty participate as mentors. Synergy exists among research mentors’ projects, and several members have ongoing collaborations. Collaborations have focused on several communicable and noncommunicable diseases, such as hantavirus pulmonary syndrome, adult T-cell leukemia, AIDS, progressive multifocal leukoencephalopathy, dengue hemorrhagic fever, WNV-associated encephalitis, avian influenza, malaria, cancer, lung disease, diabetes, and obesity.

### Student Recruitment, Selection and Curriculum Activities

The eight-month MHIRT program occurs over three semesters: Spring, Summer and Fall (Table 1). Student recruitment begins in the fall semester. MHIRT Program Directors request directors of biomedical and student research programs to provide a link to the MHIRT program on their websites. Flyers are posted on student academic boards and email information via listservs to potential applicants. Finally, former trainees are asked to assist in recruiting the best students possible.

**Table 1.** Overview of the University of Hawai‵i-MHIRT Program.

|  |
| --- |
| ● Fall Semester: Recruitment; Application processing, interviews, trainee selection |
| ● Spring Semester: Meet once a week for a 3-credit International Training in Biosciences Research course covering 8-Steps of Research with core academic-research mentors; hands on training to obtain laboratory and computer skills to conduct research at international sites |
| ● Summer Semester (May): 10 days pre-travel 1-credit onsite workshop - learn about health disparities research; continue hands on training to obtain skills to conduct research at international site, leave for international research sites |
| ● Summer Research Experience (June and July): 5-credit 8-9 weeks summer research at designated international research sites |
| ● Summer Post-Travel Workshop (August): 1-credit 10 days onsite post travel workshop - analyze data, prepare poster and oral presentations for the *E Ho`oulu Hauāmana* finale attended by faculty, staff, and family members; award presentation |
| ● Fall Semester: Continue to engage in research at the University of Hawai`i |

During November-December, trainees are selected by the Advisory Committee and members of the core academic-research mentor group after evaluating students’ essays and formal interviews. The training program takes place during the academic year beginning in the spring semester. Selected trainees register in the spring semester for 3 credits for “International Training in Biosciences Research,” for their research training and cultural competencies through topics that include 8 Steps of Research (Table 1). During this spring course, a list of possible research topics in the foreign countries are given to students to select their top three topics, and the MHIRT Program Director assigns the students their project topic.

Students first meet with their foreign mentors over scheduled videoconferencing, and they, along with the UH mentor, discuss the research being conducted at the foreign site. The foreign mentors also provide a brief overview about the country and its culture. The UH mentor then helps the trainee find, read and discuss literature on the culture of the assigned country. Based on the aforementioned literature the students propose an international cultural project to be conducted during their 8-9 weeks stay in the country.

In May, trainees participate in a 10-day workshop - (“Introduction to Biomedical Research”, 3-credits)” at the UH and then spend eight weeks in June-July at the international research site (“Research Abroad,” 5-credits). After returning in August, students participate in a 10-day workshop (“International Health Disparities”, 2-credits). Trainees attend post-travel workshops where they discuss their summer research experiences in a group setting, share their cultural projects and experiences, work with biostatisticians on data analysis, discuss research results, meet with faculty mentors, and begin preparing written reports. In the trainees-led “*E Hou‵olu Hauāmana*” program, the trainees design presentations of their research results and their cultural projects for their families, friends, and faculty. Trainees also present their results at symposia during the fall semester. Students often continue conducting research in laboratories at UH and serve as mentors for the next group of trainees.

## Results

MHIRT program evaluation and student tracking provide results on students’ academic backgrounds and performance, academic accomplishments, and attitudes related to research and a research career. MHIRT trainees were 60% women. Twenty-five percent of the trainees were Native Hawai‵ians or Pacific Islanders, 55% were other underrepresented minorities including Filipinos, and 8% were from low socio-economic backgrounds or rural areas (Table 2). Note that Filipinos are classified as underrepresented in sciences by the UH.

**Table 2.** MHIRT 2014-2019 Students’ Gender and Ethnicity.

| MHIRT<br>2014 - 2019<br>60 Students | Gender |  | Ethnicity |  |  |  |  |  |  |  | Other Criteria |  |
| --- | --- | --- | --- | --- | --- | --- | --- | --- | --- | --- | --- | --- |
|  | Women | Men | Filipino | Hawaiian | Pacific Islander | Hispanic | American Indian | White | African American | Other<sup>1</sup> | Socio- Economic Background | Rural Area |
| Count | 36 | 24 | 26 | 8 | 7 | 3 | 2 | 1 | 1 | 12 | 3 | 2 |
| % | 60 | 40 | 43 | 13 | 12 | 5 | 3.3 | 1.7 | 1.7 | 20 | 5 | 3.3 |
| <sup>1</sup>Thai, Vietnamese and Japanese; Few trainees in the White and other ethnicities categories also belonged to other criteria categories |  |  |  |  |  |  |  |  |  |  |  |  |

The mean GPA of trainees when accepted into MHIRT was 3.6 and the median GPA was 3.55. Since the program focused on infectious diseases, 77% of the trainees were from the biological sciences, and 14% were Public Health majors. Trainees from non-biological science majors were also from Psychology, Engineering, Nutrition, Economics and English.

### Student Research and Career Knowledge and Attitudes

MHIRT program evaluation systematically collected and analyzed data, documenting outcomes from students’ participation in the program. Beginning MHIRT cohort 4, we started tracking student outcomes related to research and career attitudes. Research and career related knowledge and skills were self-rated using 30 survey questions on 5-point Likert scales, (not at all - a lot). For cohort 4, the surveys were administered at the end of the program in August 2017 (data not shown). For cohorts 5 and 6, surveys were administered pre-, mid-, and post-MHIRT programs (Figure 4). These survey questionnaire items were based on evaluation metrics for undergraduate research experiences, including for minorities (Byars-Winston et al, 2016).

**FIG 4.**
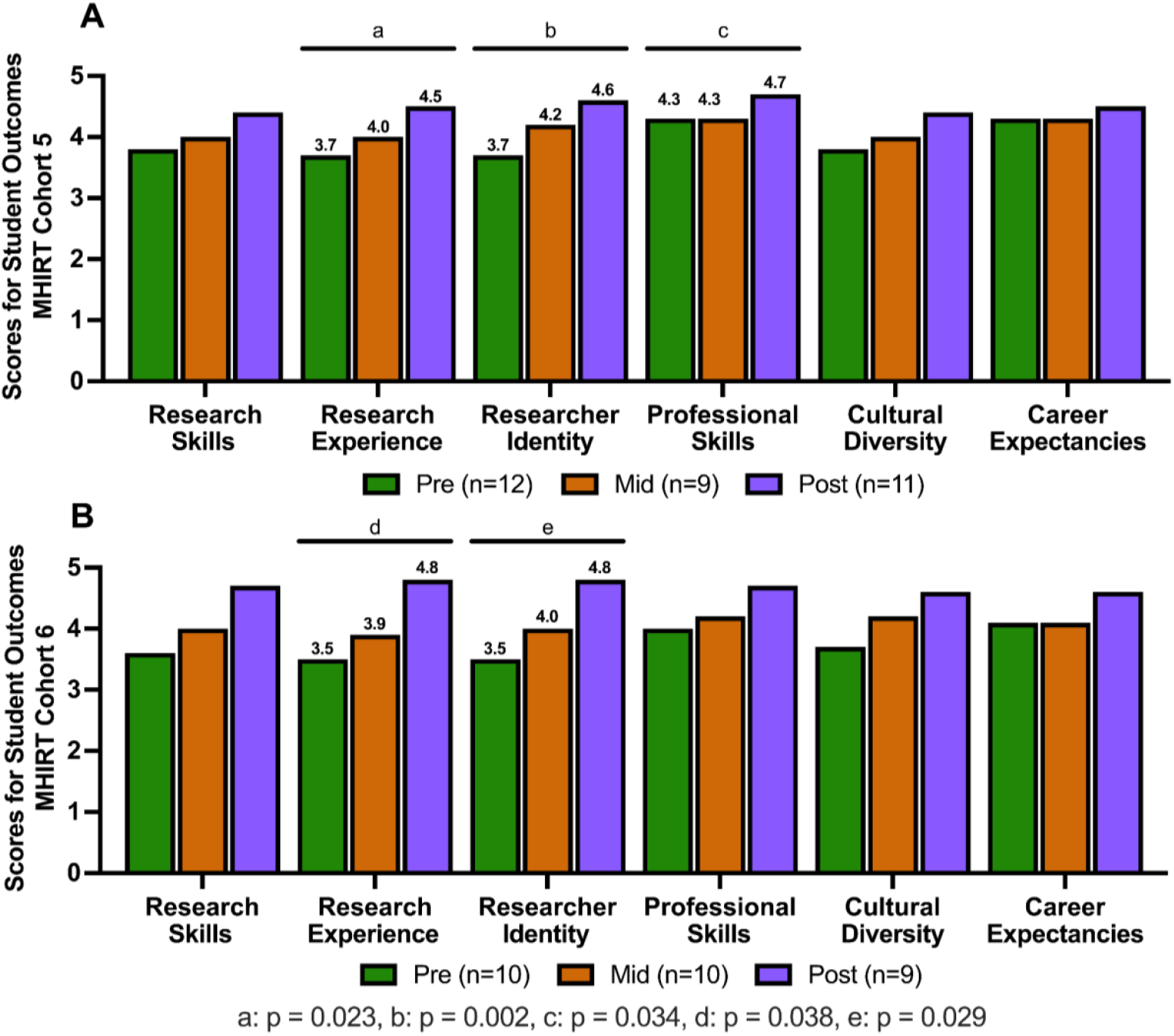
Students were administered pre-mid-post program surveys for six program outcomes measures (x-axis). Student survey outcomes for MHIRT **(A**) Cohort 5 and (**B**) Cohort 6 were measured using a 5-point Likert scales from 1 = not at all to 5 = a lot. Data was analyzed using SPSS 20.0.1.0. Kruskal-Wallis test for non-parametric data was used to evaluate group differences between pre to post surveys. ^a^p=< 0.023, ^b^p=< 0.002, ^c^p=< 0.034, ^d^**p=< 0.038, ^e^p=< 0.029.

We report students’ increases in their attitudes about research and career aspirations in cohorts 5 and 6 when students’ attitudes were tracked throughout the program (Figure 4A and 4B). For both cohorts, students demonstrated statistically significant increases on various measures. In cohort 5, there was a statistically significant increase in three categories, research experience/identity/skills, between pre- and post-evaluations. In cohort 6, statistically significant increases were also observed in two categories; research experience/identity. Other measures, e.g., cultural diversity, and research career expectancies, also increased in scores, but were not statistically significant.

Students’ attitudes were also compared at the end of the post program completion between cohorts 4, 5 and 6. There is a general trend of increase in post-program scores in each year of the program on most measures, and all post-program measures were highest in cohort 6 when compared to cohorts 4 and 5 (Figure 4 and data not shown).

At the end of the program, cohorts 5 and 6, trainees also completed (anonymously) open-ended questions related to their satisfaction with the program, what they learned, and their future plans. Students discussed their personal and professional growth because of challenges from their research experiences. When specifically asked about personal and professional skills acquired from participating in MHIRT, students identified communication and teamwork, which they felt were critical for their research and work abroad. Additional evaluation methods and qualitative results including students’ quotes are provided in the supplementary data section (Supplement 2)

### Scholarly Outcomes

As a result of the international infectious diseases training, 60 MHIRT trainees submitted 94 abstracts to local and national meetings and conferences. Further, 12 (20%) trainees published their summer research as co-authors in nine peer-reviewed journals.

### Academic and Career Outcomes

Students’ enrollment in graduate and professional programs are tracked. Of the 60 trainees who completed their undergraduate degrees, 36 (60%) enrolled in the graduate and professional programs that include MS, MD, Graduate Certificate, DO and JD. Thirty two percent were enrolled in professional health sciences programs (e.g. MPH, MD, DDS, DO), 23% were enrolled in advanced academic degree programs (GCERT, MS, PhD, post-doc) and 5% went to law school. Other alumni enrolled in and completed master’s degrees in graduate and professional (MPH, JD) schools, and others enrolled in and completed post baccalaureate programs. As of this writing, 40% of MHIRT alumni who are now working as professionals have masters, doctoral, or professional degrees. Among the academic research disciplines, 60.4% have pursued infectious disease programs.

Apart from the research projects, all students participated in cultural co-curricular activities, which provided a community context in which they conducted their research abroad. For example, i) students conducting mentored research in Pune, India joined a six-week course in yoga and meditation, ii) students presented the rich history of fabrics in Cameroon and the significance of its various designs in comparison to fabrics patterns in Hawai’i, iii) a student who dances hula in Hawai‵i participated in Thai dancing and compared it to her hula practice. They wrote about those experiences in their weekly reports, and presented them as part of their final research presentations.

## Discussion

The Department of Tropical Medicine, Medical Microbiology, and Pharmacology at the University of Hawai‵i aims to create a pipeline for underrepresented students to pursue biomedical research careers from high school to higher education including doctorate, post-doctorate and professional degrees. MHIRT is one of a few programs at TMMMP that bridges the gap in supporting the pipeline from undergraduate to professional degrees. The University of Hawai‵i MHIRT program provides a year-long classroom and experiential based research training program where undergraduate students learn about and acquire research and cultural competency, experience a hands-on research project abroad, conduct a cultural project abroad, and produce research products including oral presentations and posters on their mentored research projects. Institutional and faculty partnerships are developed through the students’ research projects.

The program evaluation and tracking strategies included both qualitative and quantitative assessments to assess research-training outcomes. Results from cohorts 5 and 6 demonstrated that students improved on measures from when they first entered the program in January to their graduation in August on the perceptions of their research identity, professional development skills, and research experiences. The comprehensive quantitative and qualitative data collected in the last two MHIRT cohorts (5 and 6) when data were available for the process evaluation indicated that MHIRT positively impacts the immediate outcomes for students’ attitudes about research and future careers, i.e., researcher identity and research experience. The experiential aspects of the program may explain the statistically significant increases in the measures that more directly impacted students’ immediate experiences. Though not significant, other measures, e.g., cultural diversity, and research career expectancies, had increased scores from pre- to post-program. Growing personally and professionally, reinforcing interest in science and research, and overall confidence and knowledge as a researcher were described and expressed by students. Furthermore, students rated all workshops very positively.

MHIRT evaluation of alumni and their career outcomes are continuously tracked by MHIRT leadership using social network platforms and once a year via email. The majority of students pursued further studies related to infectious diseases contributing to workforce development in infectious disease health research. Among the academic disciplines, 60.4% were in infectious disease programs.

## Conclusions

Our data informs global infectious diseases education and mentored research while addressing pedagogical best practices in conducting microbiology focused research. The UH-MHIRT program provides an example of a student immersion program in infectious diseases international research and country-specific cultural projects providing potential benefits for the student, and faculty and student research efforts. This may be particularly useful for programs using an international collaborative research approach to educate students to increase research knowledge and awareness about cultural diversity and sensitivity to the health issues of international populations. This is especially critical to address future outbreaks and pandemics. Most trainees in this program pursued academic graduate or professional health degrees. MHIRT has successfully trained underrepresented students to pursue biomedical infectious disease studies.

## Acknowledgements

This project was supported by a grant (T37MD008636) from the National Institute of Minority Health and Health Disparity (NIMHD), and grant (P30GM114737) from the National Institute of General Medical Sciences, National Institutes of Health (NIH) and from the UH Manoa Provost Office. We thank the mentors and staff of the following universities and organizations for their continued support of the University of Hawai‵i MHIRT trainees: University of Hawai‵i at Manoa, University of Hawai‵i at Hilo, Kapiolani Community College, Hawai‵i; Mahidol University, Chiang Mai University, Rangsit University, Thai Red Cross AIDS Research Institute, Armed Forces Research Institute of the Medical Sciences, Thailand; University of Liberia, Liberia; University of Yaounde, Centre Pasteur du Cameroon, Cameroon; Palau Community College, Palau; Lao Friends Hospital for Children, Laos; SCES Indira College of Pharmacy, Jawaharlal Nehru University, Public Health Research Institute of India, India; and University of Munich, Germany. We thank Laarni Igawa, Keeton Krause, Rennsilve Salomon, Aira Mae Corpuz, Dr. Eileen Nakano and Cori Watanabe for their dedicated assistance with the MHIRT program.

