## Supplement 1 for "Development and Implementation of a Minority Health International Infectious Diseases Research Training Program"

**Affiliations:** ^1^Department of Tropical Medicine, Medical Microbiology and Pharmacology, John A. Burns School of Medicine, University of Hawai`i at Manoa, Honolulu, Hawai`i; ^2^Department of Microbiology and Immunology, Faculty of Medicine, Mahidol University, ^3^Institute of HIV Research and Innovation, Bangkok, Thailand; ^4^Chiang Mai University, Chiang Mai, Thailand; ^5^University of Yaounde, Yaounde, Cameroon; ^6^University of Liberia, Monrovia, Liberia; ^7^Laboratory of AIDS Research and Immunology, School of Biotechnology, Jawaharlal Nehru University, New Delhi, India; ^8^ SCES’s Indira College of Pharmacy, Pune, India; ^9^Public Health Research Institute of India, Mysuru, India; ^10^Department of Native Hawai`ian Health, John A. Burns School of Medicine, University of Hawai`i at Manoa, Honolulu, Hawai`i

**Corresponding author:** *Vivek R. Nerurkar, DMLT, MSc, PhD - Professor, Department of Tropical Medicine, Medical Microbiology and Pharmacology, John A. Burns School of Medicine, University of Hawai`i at Manoa, Honolulu, Hawai`i, USA

| **Core competency** | **Learning objectives** | **Assignments** | **Date Completed** |
| --- | --- | --- | --- |
| 1. Define health disparities and health disparity research | a) Demonstrate an understanding of health disparities affecting minority populations, b) Evaluate different health disparities affecting minority populations in the country you plan to visit | - Participate in group discussions on health disparities - Attend lectures on global health topics and other health disparities issues affecting Hawaii - Research health disparities in host country and write <1,000 word essay on a specific health disparity affecting your area of research/inquiry | ☐ ____________  ☐ ____________  ☐ ____________ |
| 2. Understand what constitutes a good research question or topic | a) Define the characteristics of a good research project/question, b) be able to describe the difference between a question, hypothesis and theory | - Provide the answers to the following questions: a) list the characteristics of a good research question, b) provide an example of a good question, c) provide an bad example and tell us why it is a bad question/topic | ☐ ___________ |
| 3. Learn the Basic Steps of research and apply them to an individualized research project | a) Understand the strengths and weaknesses of laboratory-based research, b) Learn about community-based participatory research and how it can be implemented c) Demonstrate an understanding of how to conduct biomedical and behavioral research that involves grassroots communities or the lay public such as the Native Hawaiian and Pacific Islander populations | Prepare a written report detailing the study design of the project you will research this summer  Attend course discussions on the topic  • Complete an online course "Community 101" - an introductory learning module on engaging minority communities in research enterprises  • Attend the workshop lecture on CBPR and Native Hawaiian health issues.  • Participate in the discussion during the workshop lectures | ☐ ___________   ☐ ___________  ☐ ___________  ☐ ___________  ☐ ___________ |
| 4. Understand responsible conduct of research, research ethics, and cultural sensitivities | a) Understand the different research responsibilities ranging from formal regulations to good judgment and personal integrity, b) Be able to identify research misconduct and the policies involved. | • Complete the biomedical research ethics course Collaborative Institutional Training Initiative online  • Attend pre-research workshop talk on research ethics and international research  • Participate in discussion during the workshop lectures | ☐ ____________  ☐ ____________  ☐ ____________ |
| 5. Know the principles for common laboratory techniques, acquire laboratory skills needed for conducting research, and data analysis | a) Conduct basic molecular biology, immunology, cell culture techniques independently, b) Understand the different study designs and demonstrate proficiency in basic statistical data analysis | • Attend lectures on the basic principles of common molecular biology techniques, such polymerase chain reaction, DNA and RNA extraction, and flow cytometry  • Complete general lab biosafety and hazardous waste trainings  • Consult research data with the biostatistician | ☐ ____________   ☐ ____________ ☐ ____________ |
| 6. Appreciate cultural values of others, including within the MHIRT group in Hawaii and abroad | a) Be familiar with different cultural and living practices while abroad, b) Explore and highlight a cultural topic unique to the host country, c) Develop a strong professional mentor-student relationship to further research and educational development | • Attend course and workshop lectures on the different cultural norms and practices abroad  • Students write weekly reports on their observations on cultural differences and research practices  • Present a PowerPoint presentation on the cultural topic of interest | ☐ __________  ☐ __________  ☐ ____________ |
| 7. Understand the history and cultural norms of the foreign country where you will be conducting research | a) Research the history and culture of the country you plan to visit, b) Learn from your fellow MHIRT students the history and culture of the countries they plan to visit. | • Visit various websites - WHO, CDC, FBI, Homeland Security etc. and write about the history, culture, language, religion, economy, health index, etc. of the country.  • Participate in a group talk on the history & culture of the country you will visit | ☐ ___________  ☐ ___________ |
| 8. Acquire skills for working as a student member of a research team in a foreign country | a) Get to know your mentors, b) Get to know your co-workers, c) Understand basic technology you will use to conduct experiments from the point of view of your coworkers, d) Understand the local laboratory safety protocols, e) know your role in the project and lab | • Engage with your mentor and co-workers  • Plan specific activities inside and outside the workplace on weekdays and weekends. • Observe carefully experiments conducted by your co-workers | ☐ __________ ☐ ___________  ☐ ___________ |
| 9. Communicate research objectives, results and their importance to a wide audience | a) Demonstrate written proficiency in scientific writing, b) demonstrate effective oral communication skills to the public and professional audiences, c) apply critical thinking skills of scientific writing and presentations | • Conduct a comprehensive literature review on the assigned research topic  • Write a research summary of the research project  • Create an "elevator speech" that would briefly describe your research interests and MHIRT experience • Present a scientific poster at the annual JABSOM Biomedical Symposium  • Present research and cultural experiences to peers, mentors and families at the *E Ho'oulu Haumana* student presentations. | ☐ ___________  ☐ ___________ ☐ ___________  ☐ ___________  ☐ ___________ |
| 10. Know how to obtain information and apply for educational training beyond the BS/BA (e.g., MS, PhD, MD etc.) | a) Understand various academic programs offering higher education degrees, b) understand various funding agencies supporting educational needs of minority applicants | • Meet academic counselors • Seek advice from academic and research mentors • Meet financial counselors and funding agency managers  • Create a list of possible academic/research counselors and funding agencies. | ☐ ___________ ☐ ___________  ☐ ___________  ☐ ___________ |
