## Supplement 2 for "Development and Implementation of a Minority Health International Infectious Diseases Research Training Program"

**Affiliations:** ^1^Department of Tropical Medicine, Medical Microbiology and Pharmacology, John A. Burns School of Medicine, University of Hawai`i at Manoa, Honolulu, Hawai`i; ^2^Department of Microbiology and Immunology, Faculty of Medicine, Mahidol University, ^3^Institute of HIV Research and Innovation, Bangkok, Thailand; ^4^Chiang Mai University, Chiang Mai, Thailand; ^5^University of Yaounde, Yaounde, Cameroon; ^6^University of Liberia, Monrovia, Liberia; ^7^Laboratory of AIDS Research and Immunology, School of Biotechnology, Jawaharlal Nehru University, New Delhi, India; ^8^ SCES’s Indira College of Pharmacy, Pune, India; ^9^Public Health Research Institute of India, Mysuru, India; ^10^Department of Native Hawai`ian Health, John A. Burns School of Medicine, University of Hawai`i at Manoa, Honolulu, Hawai`i

**Corresponding author:** *Vivek R. Nerurkar, DMLT, MSc, PhD - Professor, Department of Tropical Medicine, Medical Microbiology and Pharmacology, John A. Burns School of Medicine, University of Hawai`i at Manoa, Honolulu, Hawai`i, USA

**Evaluation**

These results describe additional evaluation measures and outcomes of the Minority Health Research International Training (MHIRT) Program. Specifically, we collected open ended feedback about research related outcomes and satisfaction with research training topics covered in the curriculum.

Description of Survey Instrument

In 2018, student program satisfaction and career aspiration outcomes began to be evaluated throughout the eight months of the program occurring January to August. A pre-mid-post student survey asked students to rate on a 5-point Likert scale (not at all-a lot) on questions related to their perceptions of their research knowledge and skills and career interests and plans. These survey questionnaire items were based on evaluation metrics for undergraduate research experiences, including for minorities ([Byars-Winston et al, 201](https://drive.google.com/file/d/1zuXCqMjnS_6grXsyX8ItW2EZGfg8wSPq/view?usp=sharing)6).

For perceptions about their research and professional conduct students were asked questions to rate on a 5-point scale in three categories. 1) Research identity, Examples are: i) I have confidence in my ability to contribute to science, ii) I am comfortable discussing scientific ideas with others. 2) Professional skills, i) I am able to work independently, ii) I seek opportunities to explore and prepare for a career. 3) Research experience, i) I understand what everyday research is like, ii) Research frequently involves setbacks.

Open Ended Program Outcomes

Post surveys also contained open ended questions. Students provided descriptive responses about what they learned from the overall program and especially about their professional and international research experiences. The themes below validate the survey results addressing students’ increased research identity, professional skills development, and their knowledge about research experiences. Most noteworthy is that students described that their interest in pursuing research increased and/or they saw the value of research for communities.

- “I was considering only pursuing an MD career but this MHIRT experience has really opened my eyes to show me the significance of research and how it would tie in very well with an MD career.”
- “It has allowed the conversation of pursuing an MD/PhD as a future career again. I have learned that there really is no disadvantage of pursuing this career. Each one will help the other and only make you better.”
- “I think that the best thing that MHIRT did for me was push away some of my doubts about pursuing this field. I was always the average person when it came to grades and school, and never did I think that I would get these experiences and what doors I would be able to open with a degree in Biology.”
- “MHIRT helped me realize that I am serious, passionate, and want to have a career in the health field. I see myself doing it for the rest of my life and see why it is important we have researchers and physicians. Like we learned in the post-travel workshop, after researching and the development of things such as medications, we must go out into the community and work with the community to further the research.”

Students explained that this increased research interest and seeing the value of research developed because they specifically learned research and other professional skills in the program:

- “MHIRT helped me learn more research skills which I did not have before. This will help me in pursuing my health career goals because of the skills that I have gained and the experiences that I have these past months. Also, being in the MHIRT program helps me expand my professional social network as well.”
- “It has taught me important skills that can be carried into the workforce. It pushed me out of my comfort zone and has broadened my perspective on health issues affecting everyone.”
- “MHIRT helped me learn more research skills which I did not have before. This will help me in pursuing my health career goals because of the skills that I have gained and the experiences that I have these past months. Also, being in the MHIRT program helps me expand my professional social network as well.”

Other professional skills that students consistently described that they gained was improved communication skills and collaboration from participating in their research and work abroad.

- “Through this program, I was able to improve my communication skills and was also able to mentally grow not only in the lab but out of the lab as well through the experiences.”
- “I learned about how to be more patient with lab equipment/limitations and how to communicate well to others who don't speak the same language as me. Overall, communication and teamwork is especially important when you are abroad.”
- “I definitely had to learn patience and the significance of communication and teamwork…Teamwork easily translates globally. Sometimes you must take on the role of the leader and other times you must take a step back and play the support role. Each person has a strength and weakness and it is important that you are all able to work through them.”

In addition to describing the research and professional skills that they acquired, students described their personal growth qualities that were fostered in the program including knowledge of expanded future opportunities. These qualities may be especially unique to MHIRT students because for many students, this is the first time that they have gone abroad, and for some students, left the islands of Hawaii.

- “MHIRT will help me with my goals because of everything that I have found for myself. Beyond the research and literature, I have grown as a human being and developed into a stronger and more well-rounded person. Almost anyone can learn research techniques, but there is none who can teach you how to motivate yourself to create opportunities. These are the things that can be applied to any career.”
- “What I like best about MHIRT is the opportunity that it has given me to learn a new culture and valuable experiences that were life changing to me. MHIRT challenges me to be more independent in many situations.”
- “MHIRT has helped me by giving me experience working internationally. I feel as if this would help me with my health career goals as it gave me the opportunity to grow as a person and open my mind to the world around me. I think it also gave me experience working collaboratively with a diverse group of people.”

Students described that their international experiences also fostered adaptability including in living in different cultures which also resulted in their personal growth and professional skills development.

- “What I like best about MHIRT is the opportunity that it has given me to learn a new culture and valuable experiences that were life changing to me. MHIRT challenged me to be more independent in many situations.”
- “Not only did I feel challenged and responsible for the research I conducted, but I also felt as if it was my obligation to maximize my time abroad. This meant that I pushed myself to work harder and smarter each day, while also trying to immerse myself in the local culture.”
- “I liked that we were able to experience different cultures and experience conducting science in a different setting. It really allowed me to grow as a person and learn skills that will help me throughout life.”

Finally students mentioned how the program fostered their independence.

- “At the start of the program, I was nervous about my abilities. By the end of the program, I felt reassured that I can accomplish so many things independently.”
- “I would've not learned about the disparities of research in other countries. I also would have not have learned how to independently solve issues independently because everyone was really busy doing their own work.”
- “What I like best about MHIRT is the opportunity that it has given me to learn a new culture and valuable experiences that were life changing to me. MHIRT challenges me to be more independent in many situations.”

These findings provide the descriptive explanation about students’ perceptions about their experiences in the program and their program outcomes about professional and personal benefits which aligned with the quantitative evaluation survey results of the program. A student sums up how the MHIRT program provided exposure to career opportunities, strengthened research skills, and improved personal qualities: *“I feel much more confident in pursuing a career in the health field. At the start of the program, I was nervous about my abilities. By the end of the program, I felt reassured that I can accomplish so many things independently.”* Students described how the skills and experiences will help them in the future, professionally and how also they see the world:

- “It has taught me important skills that can be carried into the workforce. It pushed me out of my comfort zone and has broadened my perspective on health issues affecting everyone.”
- “I liked that we were able to experience different cultures and experience conducting science in a different setting. It really allowed me to grow as a person and learn skills that will help me throughout life.”

Program Workshop Satisfaction

After each weekly Spring workshop, students were administered online a five question survey rating of the session. According to the 17 workshops evaluated, students from both 2018 and 2019 cohorts were highly satisfied. The overall satisfaction ratings (1-5, poor-excellent) of the weekly workshops was 4.6 (SD=0.58) and 4.7 (0.68) for 2018 and 2019 cohorts, respectively.

When asked about suggestions for the MHIRT program, students identified more training in research writing and oral presentations and more hands-on interactive training. That these two areas were consistently identified indicates that many students are interested in acquiring the skills critical to become a proficient researcher.
